# Brain meta-state transitions demarcate thoughts across task contexts, exposing the mental noise of trait neuroticism

**DOI:** 10.1101/576298

**Authors:** Julie Tseng, Jordan Poppenk

## Abstract

Using new methods to capture streams of neural meta-state transitions in single participants, we characterize the psychological meaning of these regular neural events. Similar to past group-average-based analyses, participants’ individual transition streams aligned to meaningful events during movie-viewing. However, our individual-based approach also afforded observation of participants’ idiosyncratic transition timing at rest. Across these two dramatically different task contexts, transitions featured similar trait-like frequency, concurrence with activation of regions associated with spontaneous thought, and suppression by attention regions. Based on this generalization, as well as the centrality of semantics to thought, we argue transitions serve as a general, implicit neurobiological marker of new thoughts, and that their frequency therefore approximates participants’ mentation rate. Finally, to contribute convergent validity and illustrate the utility of our approach for thought dynamics, we regressed resting transition rate and movie-viewing group temporal conformity against trait neuroticism, yielding a first neural confirmation of the “mental noise” theory.

## Introduction

Research on spontaneous thought has entered the mainstream of cognitive neuroscience (Christoff et al., 2016), but methodological problems remain, including the unreliable and disruptive nature of meta-cognition (Nisbett and Wilson, 1977; Seli et al., 2018), and uncertainty about how to pinpoint a thought’s beginning and end (James, 1890). As most definitions of a thought concern its contents (with some definitions emphasizing spontaneous production; Smallwood, 2013), we propose that implicit measurement of changes in semantic content offers an interesting alternative to meta-cognition. Researchers today can readily decode neurally discernable object categories (e.g., faces and houses) from spatial patterns in participants’ functional magnetic resonance imaging (fMRI) data (Norman et al., 2006); accordingly, one could infer a new thought has arisen in a participant by observing a switch from one active category to another. One important reason this strategy has not been deployed for demarcating thoughts is the challenge of reliably distinguishing large numbers of categories; also, object categories alone may be insufficient for representing the complexity of a cognitive state. However, recent perspectives emphasize measurement of thought dynamics instead of contents (Christoff et al., 2016). What if, rather than tracking the rise and fall of particular object categories, we found a way to track semantic transitions more holistically?

Along these lines, researchers agree that time-varying changes in network interactions might signal boundaries between cognitive states (Christoff et al., 2016; Kucyi et al., 2018). However, current methods either require alignment of states across a group (e.g., by viewing the same movie stimulus), referencing known states previously visited by an individual under stimulus control (a tiny subset of the range of possible states), or focus on timescales too long to be relevant to measurement of single thoughts (with windows for representation of each cognitive state that approach a full minute long). Consequently, researchers investigating spontaneous thought have been unable to implicitly observe natural thought dynamics outside of stimulus control (e.g., using resting-state fMRI), where the timing and content of new thoughts is idiosyncratic. Arguably, implicit observation of this kind is central to any understanding of thought dynamics, as disrupting spontaneous thought for the purposes of explicitly communicating information about cognitive state itself disrupts the natural progression of states (analogous to the *observer effect* in physics).

Thus, we introduce a method to implicitly identify breaks between stable periods of brain network configuration (i.e., meta-state transitions) at a single-TR timescale and using resting-state fMRI data from single participants. We leverage movie stimuli to present a preliminary psychological validation based on the correspondence of these transitions to known semantic and perceptual features, then generalize this interpretation by showing transitions to feature various similar properties when identified within unconstrained resting-state fMRI data. Finally, we illustrate the utility of our approach for understanding thought dynamics by reporting correlations between trait neuroticism and two characterizations of neural transitions that corroborate recent personality research.

## Results

### Detecting Timepoints of Interest

We conducted our analysis on the 7T Human Connectome Project dataset, which features movie-viewing fMRI (mv-fMRI) and resting-state fMRI (rs-fMRI) data gathered from 184 participants (Van Essen et al., 2012, 2013; Ugurbil et al., 2013). We converted each fMRI run into the expression of 15 known brain networks over time (see Figure S1), then reduced its dimensionality from (*15 x time*) to (*2 x time*) using t-SNE (van der Maaten and Hinton, 2008; Billings et al., 2017). In this reduced space, epochs with similar patterns of network activity fall in proximity. We hypothesized a pattern of spatiotemporal organization reflecting progression through a series of discrete thoughts, each centred around its own semantic focal point (e.g., what one will be having for dinner) serving as an attractor. Unstable network meta-states would yield dispersion in this space, whereas an attractor would cause points to cluster, yielding a worm-like series (arising from limited drift as thoughts “evolve”; Figure 1A vs 1B). We found strong evidence for spatial contiguity with participant trajectories reliably forming into series (see Figure 1C, Figure S2, and STAR Methods).

**Figure 1.**
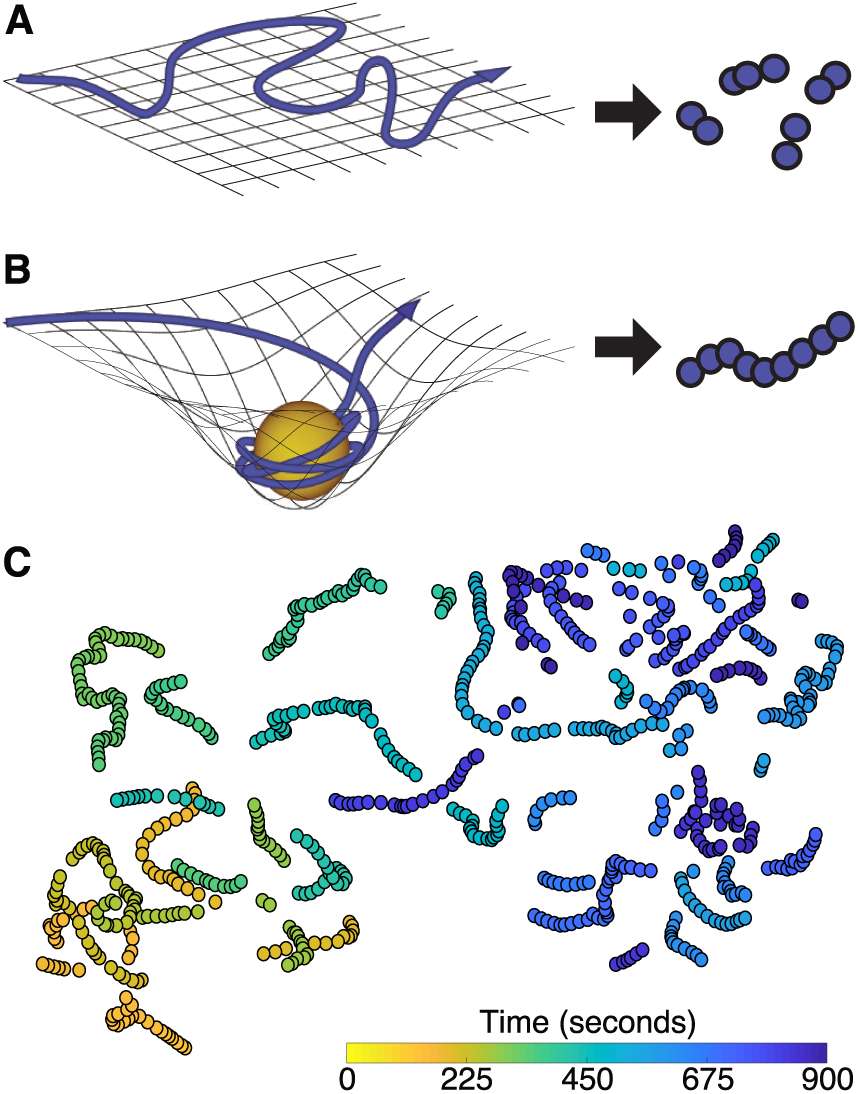
Network space representation. (**A**) Continuous, random passage through the space of possible network configurations generates fragments in t-SNE space, as opposed to (**B**) contiguous, worm-like segments when an attractor holds network configurations in relative meta-stability. (**C**) An example reduced t-SNE representation involving both segment types, as observed from one participant’s 15-minute rs-fMRI scan.

Next, we identified changes in network activity by taking the squared Mahalanobis distance (Gnanadesikan and Kettenring, 1972) between successive timepoints in t-SNE space for each fMRI run, obtaining a measure of meta-state change that we label a *step distance vector*. To stabilize the step distance vector, we repeated the dimensionality reduction and step distance vector creation process 100 times for each participant and each functional run.

Peaks within the resulting *mean* step distance vector represent prominent reconfigurations of network meta-states, thus we called them network meta-state transitions (henceforth *transitions*). For purposes of baseline comparison, we also identified local minima in the mean step distance vector, which each represent a relatively stable network meta-state (henceforth *meta-stable*) (Figure 2A).

**Figure 2.**
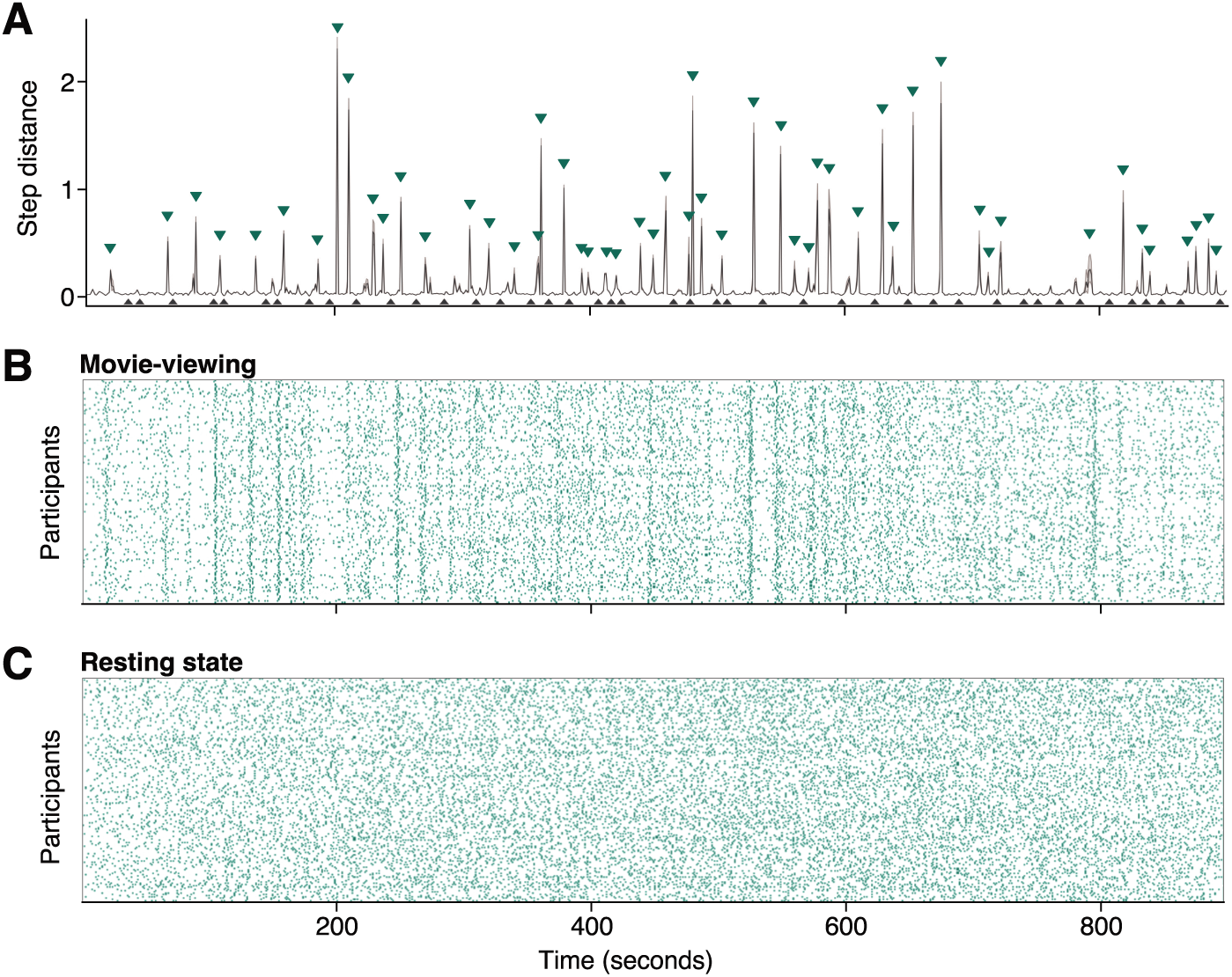
Identification of transition and meta-stable timepoints during movie-viewing and rest. (**A**) A participant’s mean step distance vector during one mv-fMRI run, with 95% CI ribbon (narrow ribbon indicates stability over t-SNE iterations). *Transition* timepoints (green triangles) and *meta-stable* timepoints (black triangles) identified by a peak finding algorithm. (**B**) All participants’ transition timepoints for the same mv-fMRI run, with many peaks overlapping those of the example participant in **A**. (**C**) All participants’ transitions for one rs-fMRI run. Alignment of peaks in **B**but not **C** reveals stimulus control over transitions.

### Leveraging Movie Data to Determine Psychological Meaning

To assess whether these discovered moments of network reorganization held psychological relevance, we validated our approach using mv-fMRI data, examining their alignment to the onset of new semantic or perceptual movie features. As a starting point, visual inspection of the set of all participant transitions during movie-viewing revealed substantial alignment in transitions relative to rest (Figure 2B, C). To quantify this, we obtained each participant’s group alignment for each resting state and movie-viewing run, which describes the correlation between the individual’s step distance vector and the corresponding median group step distance vector (i.e., conformity; see STAR Methods). We separated conformity values into runs of the same type, resulting in 723 movie conformity values and 722 rest conformity values. After feeding each set of values into a group bootstrap analysis, we found higher conformity for movie runs, mean *r* = 0.27, 95% CI: [0.26, 0.27], than for rest, mean *r* = 0.04, 95% CI: [0.03, 0.04], *r* difference = 0.23, 95% CI: [0.22, 0.25]. This finding reflects past observations of film’s unique ability to induce similar activity across participants in a wide variety of brain areas (Hasson et al., 2008; Baldassano et al., 2017; Chen et al., 2017), and provides a preliminary link between transitions and naturalistic cognition.

As an alternate means of evaluating the influence of plot progression (i.e., progression of meaning) over transitions, we attempted to predict transition alignment based on the number of narrative events in each clip. Two expert raters came to a consensus on boundaries between events (Zacks and Swallow, 2007), which we defined as timepoints where a change in the movie triggers a new semantic focal point or evolves viewer understanding of the movie narrative. We correlated the number of these events in each clip with group alignment within each participant. This yielded 184 correlation coefficients that we entered into a group bootstrap analysis. Clips with more events per minute had higher group alignment, mean *r* = 0.25, 95% CI: [0.21, 0.29].

We also directly examined the temporal correspondence of transitions with other movie features within-participant. Consensus labels of sub-events (consisting of individual actions) and cuts were obtained from two expert raters. We also obtained lower-level feature timeseries for each clip that describe semantic, visual, and amplitude change, as well as change in head motion. No other features were tested. Some features were correlated; for example, an event’s end often coincides with a cut between shots, which in turn coincides with semantic and perceptual stimulus changes. These correlations could produce a result wherein transitions appear to reflect lower-level features, but only because they peak concurrently with the onset of new cuts or events. To disentangle high- and low-level feature contributions, we “censored” epochs of lower-level feature timeseries where higher-level event boundaries co-occurred (Figure 3B). We reasoned that if lower-level features induce transitions, this should remain the case outside of censored epochs.

**Figure 3.**
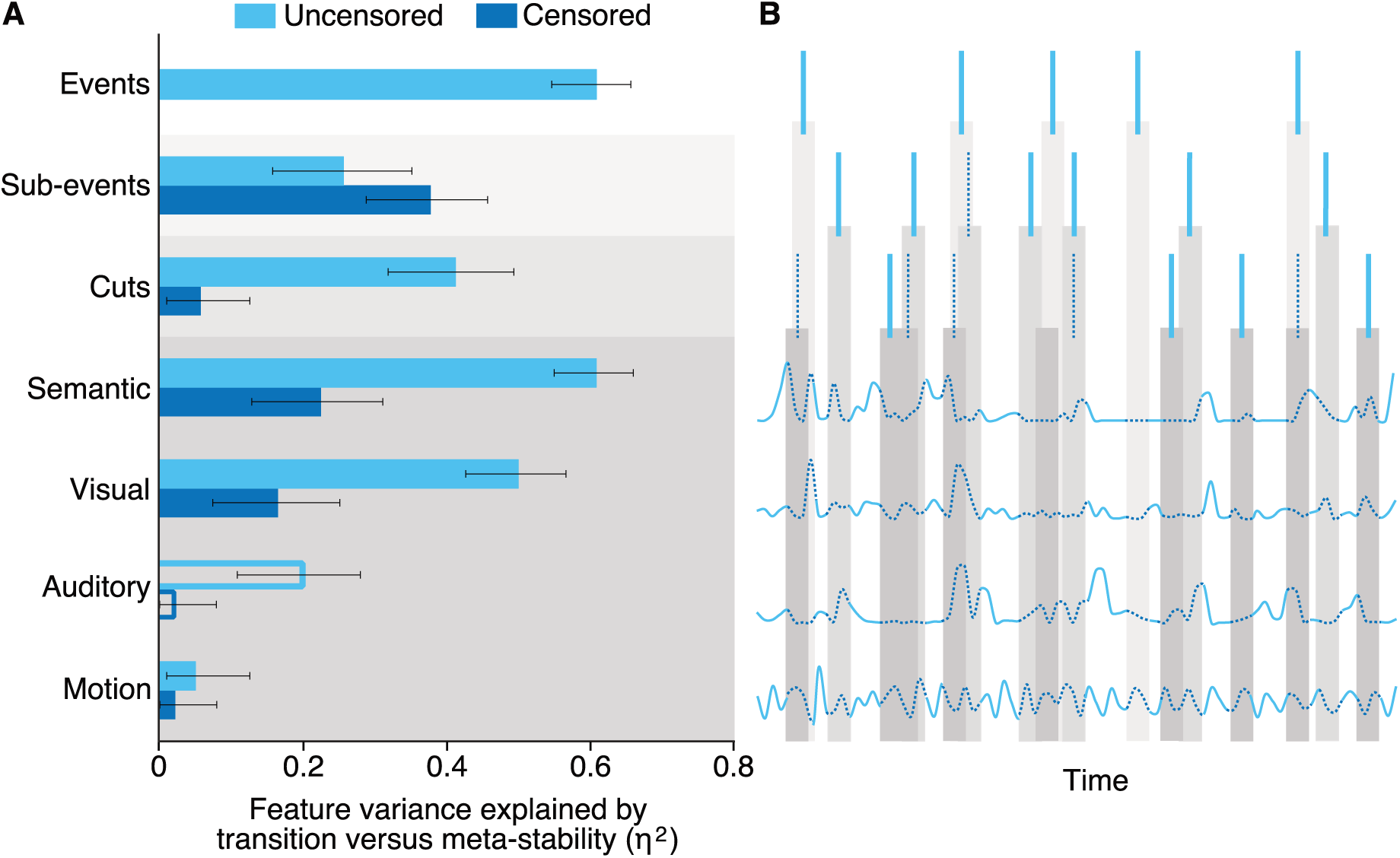
Variance in movie features explained using network meta-state transitions. (**A**) Eta-squared values describing the proportion of variance in movie features explained by alignment to transition vs. meta-stable timepoints. Error bars designate 95% confidence intervals. Filled bars denote features that are aligned to transitions, whereas empty bars denote stronger alignment to meta-stability. (**B**) Diagram illustrating censorship of lower-level feature vectors by higher-level features for greater independence of test statistics.

To determine the proportion of feature variance accounted for by transition versus meta-stable timepoints, we calculated eta-squared values for both uncensored and censored features after accounting for lag in the hemodynamic response function. Transitions (and specifically t-SNE-derived transitions over other embedding approaches or fluctuations in any of the actual 15 networks themselves; Figure S3) were strongly associated with movie features, with up to 60.8% of feature variance explained (event feature). However, with the exception of sub-events, eta-squared values for all features decreased markedly after removing timepoints that could be confounded with event boundaries. Features corresponding to non-visual perceptual change, in particular, dropped to near-zero values (Figure 3A). Thus, transitions showed a clear, albeit non-exclusive alignment to features pertaining to plot progression, with the top three predictors being event, sub-event and semantic features (see Movie S1 for illustration of this specificity).

### Extending from Feature-Rich Movies to Featureless Resting State

We interpret this co-occurrence with various movie features as evidence that meta-state transitions correspond with psychologically meaningful “mental events”. To learn whether these mental events were observable even outside of stimulus control, we interrogated possible similarities in the properties of transitions found in mv-fMRI and rs-fMRI data. First, we explored whether any consistent spatial pattern differentiated transition and meta-stable timepoints. Although transitions were defined on the basis of change in network activation, they captured both activation and deactivation of each network, so there was no circularity or bias towards the spatial characteristics of any one network.

We performed a conjunction analysis, searching for voxel activation clusters that independently distinguished transition and meta-stable timepoints for both mv-fMRI and rs-fMRI runs. We found that regardless of whether participants were engaged in movie-viewing or at rest, transitions were associated with regions implicated in spontaneous thought (Figure 4A-C, Table S1).

**Figure 4.**
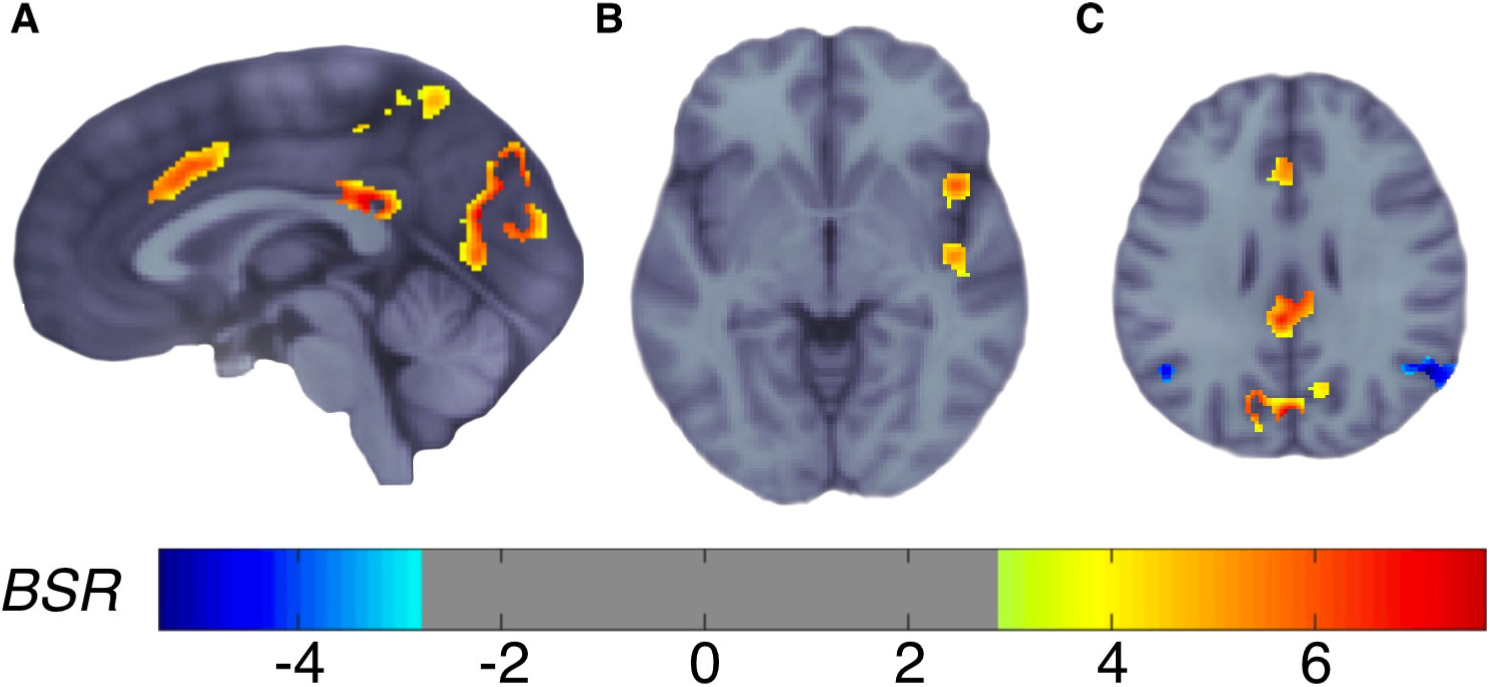
Spontaneous thought and attention regions differentiate transition from meta-stable timepoints across task and rest. (**A-C**) Images are shown in neurological orientation. (**A**) Anterior and posterior cingulate, precuneus, and visual association area were implicated in transitions. (**B**) Anterior and posterior insula were implicated in transitions. (**C**) Midline anterior, posterior cingulate, and visual association cortex were implicated in transitions; angular gyrus was implicated in meta-stability.

We also evaluated whether an individual’s *transition rate* generalized across task and rest. First, we tested its internal stability across the four rs-fMRI runs (as each was sampled in a separate scanner session). Transition rate was correlated across runs (mean *r* = 0.45, SD = 0.06), revealing it to be a trait-like characteristic that can be adequately sampled using 15 minutes of rs-fMRI data. We then averaged participant-wise transition rates across rs-fMRI runs to stabilize the trait measurement, and regressed them against the average participant-wise transition rates computed from mv-fMRI runs. Transition rate at rest was correlated to transition rate during movie-viewing, *r* = 0.60, 95% CI: [0.51, 0.70] (Figure 5), thereby further linking its properties across stimulus-driven and resting cognition.

**Figure 5.**
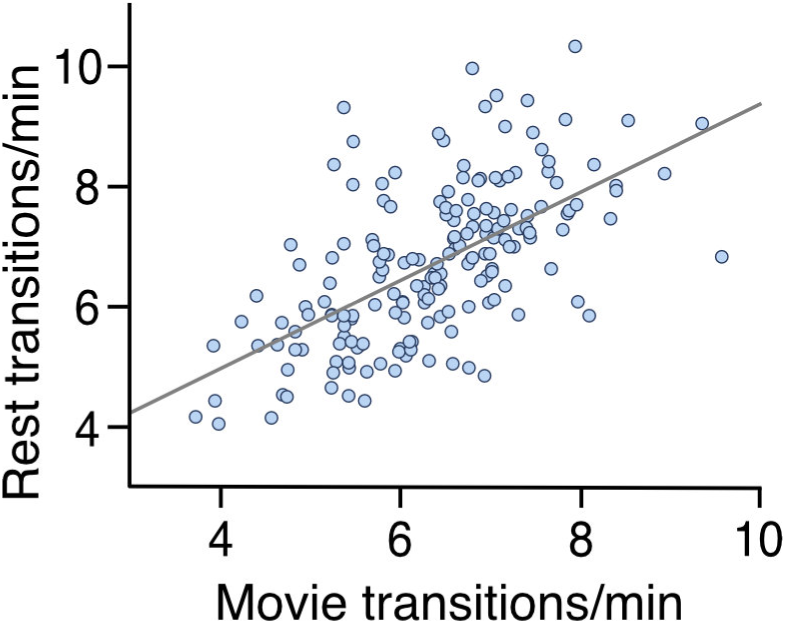
Transition structure generalizes from stimulus-driven to resting cognition. Participants’ movie-viewing transition rates correlated with their resting transition rates.

### Link Between Trait Neuroticism and Transitions

Based on the theory of “mental noise”, which suggests that individuals with higher trait neuroticism are more susceptible to distractors and lapses of attention (Robinson and Tamir, 2005), we investigated the relationship between neuroticism and our measure of transition rate while participants were at rest. Because our above analyses led us to believe that a large number of transitions corresponded to a large number of thoughts, we interpreted transition rate as analogous to mentation rate, and predicted that the higher mental noise associated with neuroticism would correspond to a higher rate of transitions. Higher levels of trait neuroticism were associated with higher resting transition rates, *r* = 0.15, *p* = .027 (one-tailed). As a further test, we hypothesized that mental noise would correspond to more idiosyncratic transitions during movie-viewing, yielding transition timing in high-neuroticism participants that conformed less to that of other participants (i.e., less alignment between an individual’s stream of transitions and the group). We note that this test drew on a different source of data than our first analysis (mv-fMRI rather than rs-fMRI). Consistent with our hypothesis, higher trait neuroticism was associated with lower conformity during movie-viewing, *r* = −0.18, *p* = .008 (one-tailed).

## Discussion

To summarize, our method characterizes a neurocognitive landmark based on transitions between brain network meta-states. Our approach is distinctive for its ability to observe fine-grained thought dynamics, and applicability to single individuals being scanned in any task context, with or without prior knowledge about the complexity of brain data under interrogation or the meta-states to be visited. As validation, we found transitions to be responsive to events in movie stimuli, and found features of transitions to be both trait-like and generalizable across task-based and task-free imaging contexts. Finally, we established construct validity of transitions by showing the frequency of their occurrence within resting-state fMRI data and lack of alignment to the group within movie-viewing fMRI data to predict higher trait neuroticism, as predicted by prevailing (behaviourally-constructed) theory in personality psychology.

A thought is grounded in its contents. Therefore, our findings that meta-state transitions during movie-viewing best predicted event boundaries and onset of new semantic information support the interpretation that meta-state transitions align with the changes in semantic content across one’s thoughts. These results complement those of Baldassano and colleagues (2017), who used a Hidden Markov Model (HMM) approach on specific neural structures to find shifts in movie-viewing brain activity which co-occurred with event boundaries. However, our method differs by its reliance on a specific and observable neural phenomenon (whole-brain network transitions), its applicability on an individual basis, and its potential capacity to detect semantic change in uncharacterized task environments.

The regions consistently associated with transitions across task contexts have been previously associated with spontaneous thought. In particular, anterior cingulate and insular cortices are considered members of a salience network that shifts attention to novel external and internal events (Seeley et al., 2007); the posterior cingulate cortex is a key region in the default mode network that is more active during task-unrelated than task-related thought (Leech and Sharp, 2013; Smallwood et al., 2016). By contrast, we found that angular gyrus activation, which has been associated with both event representation (Seghier, 2012) and sustained attention (Singh-Curry and Husain, 2009), predicted meta-*stability,* perhaps reflecting less transitions while participants focally contemplated an event.

Transition rate was correlated within runs of the same type, revealing it to be a trait-like characteristic that can be adequately sampled using 15 minutes of rs-fMRI data. This result builds on recent findings of trait-like properties of low-frequency chronnectome characteristics (Calhoun et al., 2014; Liu et al., 2018) by showing that higher-frequency network reconfigurations relevant to rapid, thought-like fluctuations in cognitive states are also trait-like. Furthermore, the average transition rate during movie-viewing was strongly related to average transition rate at rest. This relationship may be explained by the finding that participants segment commercial films and naturalistic clips with similar fractal structure (Blau, Petrusz and Carello, 2013), suggesting commonality between mentation during movie-viewing and real experiences. Film theorists lend further support to this finding, positing that narrative films and the techniques associated to creating them are not an attempt to reproduce reality, but are optimized based on real perception to control viewer’s attention and mental state (Shimamura, 2013).

Our method yielded a neural measurement of mentation rate that we used to investigate how neuroticism relates to thought dynamics. In particular, the mental noise hypothesis proposes that individuals with high neuroticism are more susceptible to intrusive thoughts and distraction by irrelevant information (Robinson and Tamir, 2005; Perkins et al., 2015; Klein and Robinson, 2019), presumably yielding more frequent and less predictable changes in cognitive state. Consistent with this proposal, we found trait neuroticism to predict higher transition rates (i.e., mentation rate) during rest, as well as lower conformity of transition temporal structure to that of the group during movie-viewing. We interpret these two observations as reflecting increased mental noise in individuals with high trait-neuroticism, which in turn supports the construct validity of transitions as a measurement of thought dynamics in both rs- and mv-fMRI. It also serves as a first neural confirmation that neuroticism indeed entails a “noisier mind”.

Taking these observations together, we argue that neural meta-state transitions can serve as an implicit biological marker of new thoughts. Although our analysis has focused on movie and resting fMRI data from healthy adults, our methods are applicable to a wide range of tasks, populations, and even case studies. Overall, by lending a level of validity and reliability unavailable using past introspective approaches to measuring thought dynamics, our approach makes new questions accessible: for example, how many thoughts do we experience each waking day? Extrapolating from a median transition rate across movie-viewing and rest of 6.5 transitions / min, and a recommended sleep time of 8 hours, one could estimate over six thousand daily thoughts for healthy adults of a young-adult demographic similar to the one used in our analysis. Although further interrogation of meta-state transitions would be needed to employ such a measure with confidence, availability of a tentative answer further highlights how the current approach may be fruitful in advancing how we think about thought.

## Acknowledgments

We thank Y. Lu for help with event, sub-event, and cut segmentations, S. Smith for manual hippocampal segmentations, and K. Norman, M. Sabbagh and G. Blohm for helpful comments. This research was funded by Natural Sciences & Engineering Research Council Discovery Grant 03637 (J.P.), which also supported J.T. Infrastructure funding was provided by the Canada Foundation for Innovation – John R. Evans Leaders Fund (J.P.). and a Queen’s University Research Initiation Grant to J.P., who was supported by the Canada Research Chairs program. Data were provided [in part] by the Human Connectome Project, WU-Minn Consortium (Principal Investigators: David Van Essen and Kamil Ugurbil; 1U54MH091657) funded by the 16 NIH Institutes and Centers that support the NIH Blueprint for Neuroscience Research; and by the McDonnell Center for Systems Neuroscience at Washington University.

## Author contributions

Conceptualization, J.T. and J.P.; Methodology, J.T. and J.P.; Software, J.T. and J.P.; Formal Analysis, J.T. and J.P.; Resources, J.P.; Data Curation, J.T. and J.P.; Writing – Original Draft, J.T. and J.P.; Writing – Review & Editing, J.T. and J.P.; Visualization, J.T. and J.P.; Funding Acquisition, J.P.

## Methods

### Key Resources

Our analysis relied upon algorithms implemented in MATLAB (v.2017a), FSL (v.5.0.10), and Datavyu (v.1.3.7).

### Lead Contact and Materials Availability

Further information and requests for resources should be directed to and will be fulfilled by the Lead Contact, Jordan Poppenk.

### Experimental Model and Subject Details

#### Human Connectome Project dataset

Neuroanatomical and functional data were collected by the WU-Minn Human Connectome Project (HCP) consortium (Van Essen et al., 2013). In a prior analysis, a 3T resting state fMRI (rs-fMRI) dataset of 1003 participants (age *M* = 28.7 years, *SD* = 3.7 years; 534 female) was used by the HCP group to generate spatial maps of typical brain networks that may be found in rs-fMRI participants through a process involving group-PCA and group-ICA (Beckmann et al., 2009; Nickerson et al., 2017). As this dataset was the source of these maps, we refer to it as the *training* dataset. Our analyses involved applying these maps to a *testing* HCP dataset (7T) that contained both mv-fMRI and rs-fMRI scans, and that was the subject of all of our own analyses. Although this 7T dataset is described elsewhere (Van Essen et al., 2012, 2013; Glasser et al., 2013; Ugurbil et al., 2013), briefly, it consists of fMRI scans from 184 participants (age *M* = 29.4 years, *SD* = 3.4 years; 112 female). Each participant underwent four 15-minute mv-fMRI and four 15-minute rs-fMRI runs; functional images were acquired using a multiband gradient echo-planar imaging (EPI) pulse sequence (TR 1000 ms, TE 22.2 ms, flip angle 45°, multiband factor 5, whole-brain coverage 85 slices of 1.6 mm thickness, in-plane resolution 1.6 x 1.6 mm^2^, FOV 208 x 208 mm^2^) (Feinberg et al., 2010; Moeller et al., 2010; Setsompop et al., 2012; Xu et al., 2012). During each movie run, participants watched three or four movie clips interspersed with 20-second rest periods as well as an 84-second validation clip repeated at the end of each run (due to its repetition, we did not include this clip in our analyses).

In addition, high-resolution T1-weighted and T2-weighted scans were gathered (TR 2400 ms and 3200 ms, TE 2.14 ms and 565 ms, flip angle 8° and variable, 0.7 mm thickness, in-plane resolution 0.7 x 0.7 mm^2^, FOV 224 x 224 mm^2^) for purposes of group anatomical alignment. Data collection was approved by the Washington University institutional review board (Van Essen et al., 2012), and performed by the HCP consortium, which also gathered informed consent from all participants at the time of data acquisition. Access to these datasets was granted by the HCP consortium, and acknowledged by the Health Sciences Research Ethics Board at Queen’s University. No participants were excluded from analysis.

### Method Details

#### Step 1: Timepoint Discovery

##### Transforming functional data to 15-network representation

We mapped the 15 spatial maps resulting from the training set group-ICA decomposition onto each 7T participant’s resting state and movie-viewing data using FSL’s dual regression function (Beckmann et al., 2009; Nickerson et al., 2017), a method in which known spatial configurations are regressed against new data to transform 4D functional data into a set of timeseries (one per spatial map, Figure S1A).

**Figure S1.**
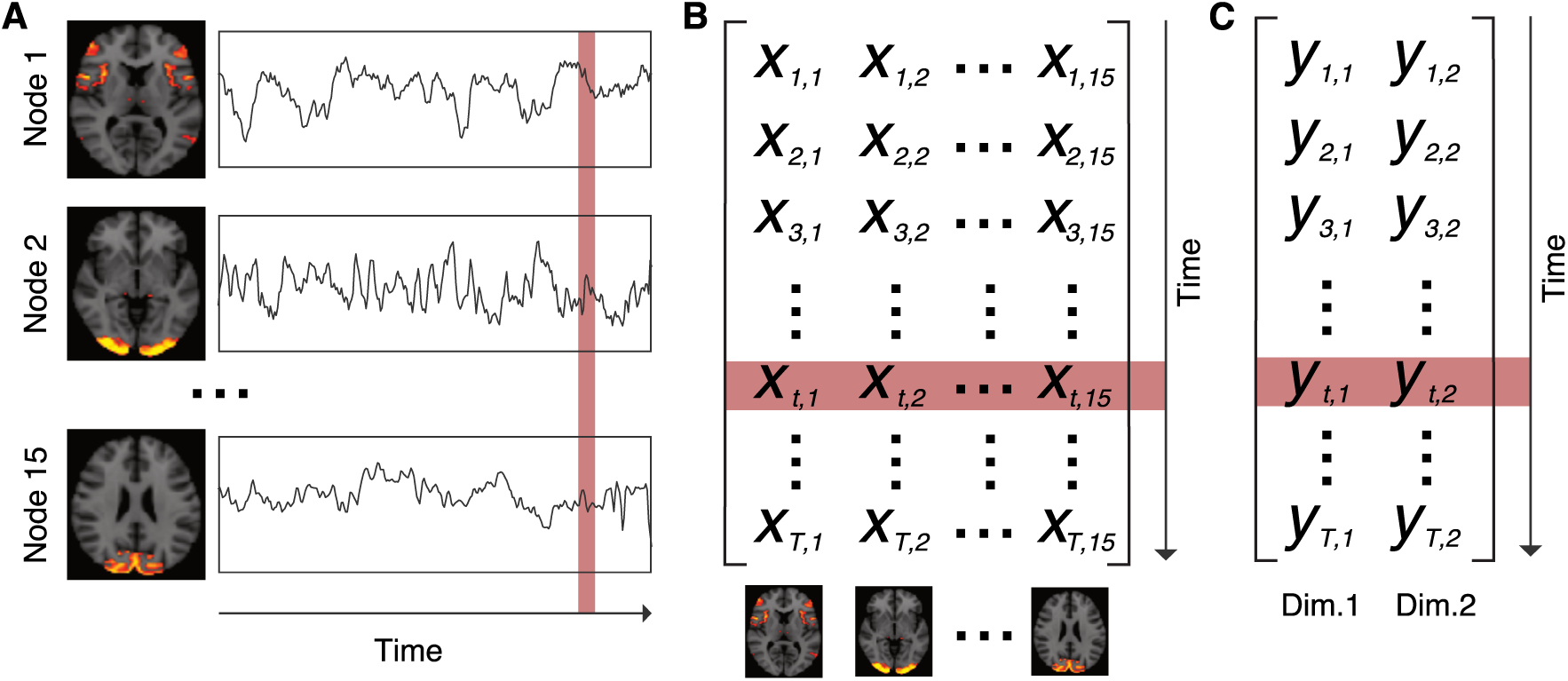
Expression of brain network activity. (**a**) Each fMRI run of each participant was characterized as the activation of 15 brain networks over time. (**b**) Representation of A as a two-dimensional matrix of (time x network). (**c**) Reduced representation obtained after applying t-SNE algorithm. Related to: Figure 1C.

Although larger network set sizes were available from the HCP group (ranging from 15-to 300-brain-network solutions), we observed that set size had little material impact on trajectory estimates, and therefore selected the simplest available (15-network) solution. To increase signal-to-noise ratio and to dampen short timescale network fluctuations, we temporally smoothed each resulting timeseries with a moving average filter (span = 5 s). This procedure yielded a smoothed timeseries for each brain network reflecting that network’s activation over time. We combined these timeseries to create separate (time x network) matrices for all four mv-fMRI and all four rs-fMRI runs for each participant (Figure S1B).

##### Transforming network representation to the lower-dimensional space of network meta-states

In preparation for using the Mahalanobis distance metric on the data, we applied the t-distributed stochastic neighbour embedding (t-SNE) algorithm to reduce the dimensionality of each matrix from 15 dimensions to 2 dimensions at the default perplexity setting of 30 (Figure 1C, Figure S1C; van der Maaten and Hinton, 2008). We found the perplexity setting had little impact on our analysis, and therefore selected what is regarded as a moderate (and default) value.

Notably, Miller and colleagues (2016) used a somewhat similar approach, creating a low-dimensional representation by first defining the space of possible meta-states as a discrete 5-dimensional state space, with each dimension representing a distinct group temporal ICA component derived from participants’ functional data (i.e., connectivity patterns). Whole brain activity was expressed as a weighted combination of these components over time. To map each timepoint onto their meta-state space, they then discretized each weight at each timepoint according to its signed quartile. In contrast, our approach relies on dimensionality reduction algorithms to discover changes in meta-state directly from the continuous-valued 15-brain-network representation. We selected this approach because it affords flexibility in the designation of each meta-state by mapping each one onto a continuous two-dimensional space instead of a discrete 5-dimensional state space. Just as the Greek philosopher Heraclitus noted, “No man ever steps in the same river twice, for it’s not the same river and he’s not the same man”, our approach is aligned to the very likely possibility that meta-states are continually evolving. Our approach also differs by drawing on published reference networks derived from a static group of 1003 participants (i.e., the training dataset described above; Beckmann et al., 2009; Van Essen et al., 2013; Nickerson et al., 2017), rather than a set of networks derived from the specific dataset under interrogation.

##### Detecting network meta-state transitions

To derive from our t-SNE representation a global measure sensitive to changes in network meta-state, we computed the Mahalanobis distance in position within this low-dimensionality t-SNE space across subsequent timepoints. The resulting *step distance vector* for each fMRI run of each participant should peak at points in the timeseries where shifts in network meta-state occur. To address potentially divergent results across repeated t-SNE algorithm runs, we repeated the dimensionality reduction and step distance vector creation process 100 times for each participant and each functional run. Then, we took the mean across the 100 step distance vectors for that run (Figure 2A). However, even 95% confidence intervals were tightly constrained.

We applied a peak finding algorithm on each mean step distance vector to identify *transition* timepoints at which the step distance satisfied a minimum peak prominence threshold of 0.06, the value at which approximately 80% of all step distance values fell under the 5^th^ percentile transition-associated step distance value. Setting a prominence value rather than applying a high pass filter allows the algorithm to consider step distances in the neighbourhood surrounding the peak being evaluated and results in more robust transition selection. To find *meta-stable* timepoints, we inverted the signal and specified a minimum peak width of 10; this parameter ensured that timepoints would only be identified within persistently meta-stable periods. One example of a participant’s identified transitions and meta-stable timepoints are shown in Figure 2A with green and black triangles, respectively. Under these parameters, the median step distance value associated with transition timepoints was 0.48 (5^th^ and 95^th^ percentile: [0.09, 1.79]). The median exceeds 94% of all values, while the lower bound exceeds about 80% of all values (Figure S2B). By contrast, the median step distance associated with meta-stable timepoints was 0.02 (5^th^ and 95^th^ percentile: [0.01, 0.03]), falling below about 94% of all values.

##### Quantifying contiguity in the network meta-state space

We examined the degree of contiguity in the t-SNE representations by comparing the standardized step distance distribution to an analogous distribution based on Gaussian noise. First, we generated simulated noise data, where each simulation consisted of 900 pairs of values taken at random from a normal distribution with the mean and standard deviation of a random participant’s movie or rest t-SNE representation (Figure S2D). We then calculated a step distance vector for this noise sample. The distribution of real data (described in Figure S2B) was comprised of 1,311,791 values collapsed across participants and all available functional runs, where each run had 900 to 921 timepoints. Thus, we repeated the noise sampling procedure 1,460 times (the number of actual values divided by 900), generating a similar number of simulated values (Figure S2E).

**Figure S2.**
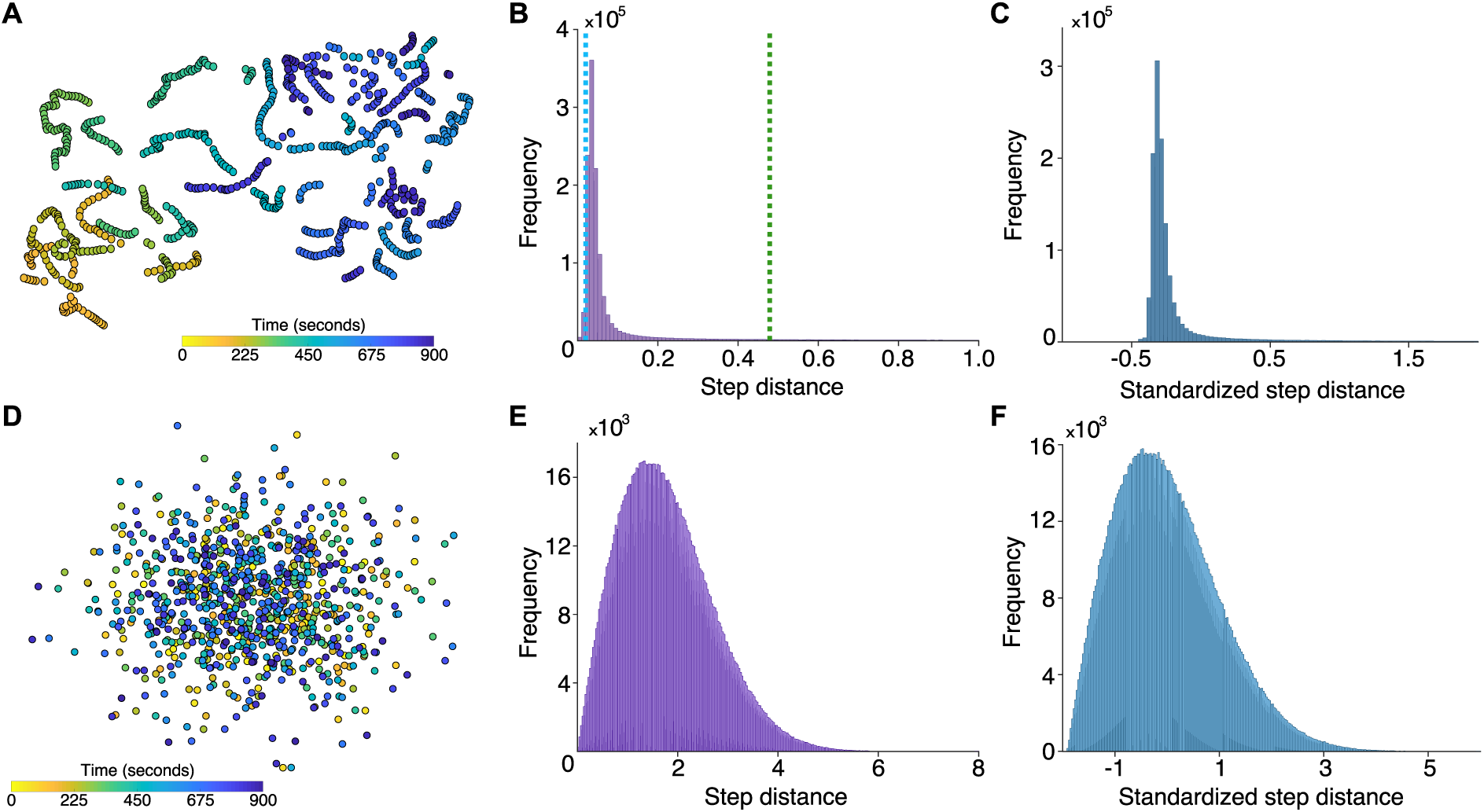
Distribution of step distance values. (**a**) An example reduced t-SNE representation involving both segment types, as observed from a 15-minute rs-fMRI scan of one participant. (**b**) A histogram depicts the mean step distances within 7T participants’ four rs-fMRI and four mv-fMRI runs. Approximately 94% of the observed values fell below the median *transition* distance (green dashed line), and 94% above the median *meta-stable* distance (blue dashed line). (**c**) A histogram depicts the standardized step distance values. Note that the values are mostly distributed close to zero, but feature a heavily skewed tail. (**d**) An example of noise simulated by sampling pairs from a normal distribution with characteristics mirroring actual reduced t-SNE representations. (**e**) A histogram analogous to the one in panel B that depicts the step distances across simulated noise samples. (**f**) A histogram depicts the standardized step distance values of noise samples. Note that relative to the analogous real data depicted in C, the simulated noise values are not as tightly distributed around zero, and the y-axis scale is two orders of magnitude less than the corresponding distribution drawn from real data in C. Related to: Figure 1C, Figure 2A.

To better compare the dispersion in the actual data against the dispersion in the noise samples, we standardized both distributions (Figure S2C, F). Standardized step distance values for real data were predominantly dispersed closely around zero, whereas the noise standardized values were dispersed over a wider range. Also, the standardized step distance distribution for real data had a skewness of 5.0 and a kurtosis of 34.3, reflecting a long tail of extreme values, whereas the standardized noise distribution’s skewness and kurtosis metrics indicated a fairly normal distribution (skewness of 0.6 and kurtosis of 3.2). Together, these distinctions help to formalize the temporal contiguity of real data that is visually apparent in Figure S2A.

#### Step 2: Psychological relevance

##### Movie feature vectors

To link our identified neural transitions to participants’ experience, we investigated how transitions mapped onto well-characterized movie features (events, sub-events, cuts, semantic, visual, and auditory), as well as each participant’s recorded head motion during the scan. Sub-events describe directly what an actor or multiple actors are doing or saying (“he is walking down the stairs”, “they are riding in a car”), or they describe the motion of important objects (“a meteorite strikes the ground”, “a car drives down the street”). We defined an event as a meaningful cluster of sub-events that describes a larger, overarching goal achieved by the sum of its parts (i.e., fine-grained and coarse-grained events; Zacks and Swallow, 2007). Cuts referred to boundaries between two separate camera shots. Two raters used a video coding tool (Datavyu Team, 2014) to independently identify boundaries demarcating events and sub-events, then met to discuss any differences in their ratings and achieve consensus event boundary timepoints. Using this approach, raters created consensus event and sub-event boundary segmentations for all four movie runs (14 clips). Only one rater logged cuts, as the objective nature of camera positions left little to discussion. Across the 14 clips found during the four movie runs, raters identified an average of 7.5 events, 37.3 sub-events, and 46.2 cuts per clip. A binarized timeseries was created for each logged feature, with onsets allocated to the nearest 1 s time bin (corresponding to the 1000 ms TR during which each fMRI volume was gathered).

The HCP group supplied two types of feature labels for the movie stimuli: semantic-category labels that described the high-level semantic features contained in each one-second epoch of the film (Huth et al., 2012), and motion-energy labels that described low-level structural features in the same epochs (Nishimoto et al., 2011). There were 859 semantic features and 2031 motion energy channels that expressed changes in the semantic content and motion-energy of each epoch, respectively. By summing across all semantic features and taking its derivative, we obtained a measure of overall magnitude of change in semantic content at each epoch. Similarly, we summed across all motion energy channels and took its derivative to obtain a measure of overall magnitude of change in perceptual features at each epoch. We also took the absolute value of the auditory amplitude vector derivative for each movie run as a measure of magnitude of change in volume. Finally, to rule out the possibility that transitions are a motion artifact, we obtained, for each participant, relative root-mean-square (RMS) change in head position (Jenkinson, 1999), corresponding to a vector of head movement over time.

##### Disentangling event boundaries and movie features

Whereas the feature vectors obtained above were correlated, we derived from them a set of independent feature vectors by “censoring” epochs where event and sub-event boundaries, as well as cuts were present. This was done by dropping values in a 3-second window around feature boundaries and ensured that apparent effects in lower-level features would not be explained by correlation to higher-level features. Event boundaries censored all other vectors; sub-event boundaries censored all vectors other than events; and cuts censored all vectors other than events and sub-events (Figure 3B). To prevent spurious effects related to clip onset within movie runs, we also censored the first six seconds of each clip for both the uncensored and censored feature vectors.

##### Movie and movement features at transition versus meta-stable timepoints

For each transition and meta-stable point found in the mv-fMRI data, we next computed the average level of each feature accounting for hemodynamic response function (HRF) lag (working backwards based on the canonical HRF (Aguirre, Zarahn and D’Esposito, 1998), sampling from a feature window 3-6 s prior to each transition and meta-stable point). We then averaged across all onsets of the same type for each participant. Thus, each participant ultimately had two values for each feature vector: one representing the average feature vector value at a transition timepoint, and an analogous value at a meta-stable timepoint.

We ran a t-test comparing these two values across participants for each feature, and used each resulting t-statistic to compute the proportion of variance that was explained in the feature vector (i.e., eta-squared) by the presence of a transition. Next, we used a non-parametric bootstrapping analysis to obtain a 95% CI for each feature (Bollen and Stine, 1990). Using participants’ transition-baseline feature value difference as input data, this approach constructs a sampling distribution of the mean by resampling 1000 times with replacement across participants. Then, the t-statistics corresponding to the upper and lower bounds of each CI were converted to eta-squared values (Figure 3A).

##### Performance of alternative embedding approaches

Having identified strong prediction of various features, we next sought to determine how crucial our specific embedding approach (i.e., t-SNE) was to identifying transitions that mapped strongly onto movie features. To this end, we derived four additional sets of transition and meta-stable timepoints: one from the unreduced (time x network) representations of each participant’s mv-fMRI data (Figure 1B), a second and third set from a 2-dimensional representation of their mv-fMRI data obtained through principal components analysis (PCA) and independent components analysis (FastICA), and a fourth from the method described by Miller and colleagues (2016).

For Miller and colleagues’ method, we began by regressing the spatial ICA maps corresponding to the 50-network decomposition into the functional data to obtain a 50-network representation of brain activity over time and match the dimensionality of their initial input data. Then, we calculated the pairwise correlations between 44 s time-window of activity in each network. This process was repeated for the entire timeseries by sliding the time-window in increments of 1 s. As a result, a given participant’s brain activity at a time-window was expressed as the set of pairwise correlations between each of the networks (i.e., a connectivity pattern). Then, we applied FastICA to obtain a 5-dimensional representation of activity over time, with each of the 5 components representing a particular connectivity pattern. After discretizing the weights of each component at each time-window according to its signed quartile, the resulting 5-dimensional representations were used to derive corresponding step distance vectors, as well as transition and meta-stable timepoints.

Specifically, for each embedding (15-dimensional for the unreduced timeseries, 2-dimensional for the PCA and ICA approaches, and 5-dimensional for Miller and colleagues’ method), we calculated the Mahalanobis distance across subsequent timepoints and applied the peak finding algorithm to identify transition timepoints; the minimum peak prominence threshold selection strategy and peak finding parameters for meta-stable timepoints remained unchanged. As above, we used these three additional sets of transition and meta-stable timepoints to obtain the proportion of variance that was explained in the uncensored feature vectors by the presence of a transition for each embedding approach (Figure 3).

Transitions derived directly from the 15-dimensional (time x network) representations predicted a moderate proportion of variance in events, but not for lower-level event-based features such as sub-events or cuts. PCA, ICA, and Miller’s method’s transitions performed worse than t-SNE transitions for all feature categories. We interpret the strong performance of t-SNE in predicting features (especially semantic features) as reflecting its unique ability to distill important local and global structure from the data.

Consistent with this idea, as dimensionality increases, distance metrics are understood to lose their usefulness as the distances to the nearest and furthest point from any reference point approach equality (Beyer et al., 1999). Zimek and colleagues further showed that if the dimensions are correlated rather than independent and identically distributed, then considering subsets of dimensions can improve the performance of distance metrics such as Euclidean distance (Zimek, Schubert and Kriegel, 2012). This likely explains the poor unreduced feature prediction in our analysis, and supports the usage of a dimensionality reduction algorithm, as the spatial networks in the 15-dimensional (time x network) representations that we submitted to analysis were not independent from each other. Substituting Mahalanobis for Euclidean distance can additionally account for residual correlations between reduced dimensions (Gnanadesikan and Kettenring, 1972).

In spite of this, in our analysis, poor PCA and ICA performance signalled that the *kind* of dimensionality reduction approach also matters. PCA and ICA are linear methods that seek to preserve global structure, whereas t-SNE can balance a trade-off between local and global structure. Upon exploring the variable loadings for PCA results, we observed that the two principal components were weighted heavily towards visual networks that likely explained substantial global variance, but at the cost of sensitivity to changes in networks related to higher-level conceptual processing. As a result, PCA-based transitions performed comparably to unreduced transitions in the visual category and correlated categories (e.g., auditory), but worse in semantic categories. The increased restrictiveness that results from the additional requirement of statistical independence imposed for ICA components may explain why it performs even worse than PCA.

**Figure S3.**
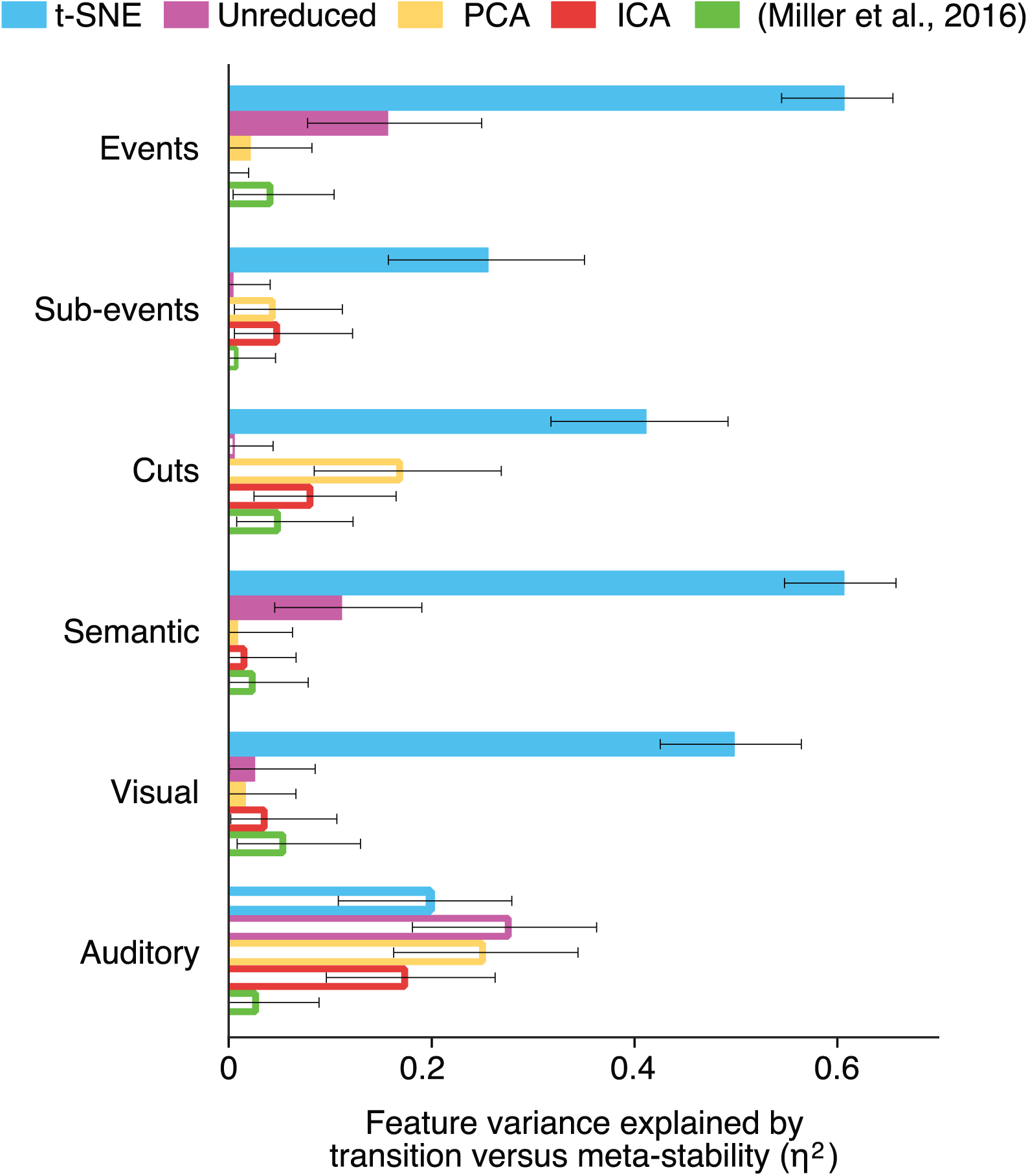
Impact of embedding approach on transitions’ predictive power for movie features. Eta-squared values describing the proportion of variance in uncensored movie features explained by alignment to transition vs. meta-stable timepoints. Filled bars denote features that are aligned to transitions, whereas empty bars denote stronger alignment to meta-stability. Colours indicate the state space from which Mahalanobis distances were calculated to identify transition and meta-stable timepoints. Related to: Figure 3.

Transitions derived from Miller and colleagues’ (2016) method were not associated with feature changes. As we are looking for moment-to-moment changes in network reconfiguration, one possible explanation is that the size of the time-window used to correlate across the 50 networks (44 seconds) was too long to be sensitive to feature-related changes at a certain timepoint. Thus, we tested the smallest time-window that could be evaluated (3 seconds), but still found that no features were associated with transition relative to meta-stable timepoints (all *p*s > .28). A 2 second time-window is impossible to evaluate as the pairwise correlation values are −1 or +1, reflecting the linear relationship between two points. Furthermore, reducing dimensionality by distilling the 50-network pairwise correlations to 5 connectivity patterns may not provide enough flexibility to capture the different mental states linked to movie-viewing.

##### Mapping between fluctuations in particular network nodes and movie features

We regressed each movie feature timeseries on each participant’s 15-network node timeseries to investigate the predictive power of individual networks. This yielded 184 adjusted R-squared values for each of the 6 movie features. The highest mean adjusted R-squared value was for the semantic feature (mean R^2^ = 0.0033), signalling that individual networks were not meaningful predictors of any feature.

##### Influence of event structure on degree of transition group alignment (conformity)

Visual inspection revealed considerable structure in transitions across participants during movie-viewing (Figure 2B), complementing prior findings of local coordination of brain activity as participants watch well-made films. To formalize this observation, we tested for higher group alignment in movie runs than rest runs. For each participant and each run, we took the Fisher transformation of the correlation between the log of their step distance vector and the log of the median group signal (excluding the step distance vector of the participant in question). To avoid potential effects resulting from the onset and offset of the run, we excluded the first and last five epochs of the step distance vectors. Then, we compared all group alignment values from the four mv-fMRI runs against zero. The same bootstrapping analysis was carried out for all group alignment values for the four rs-fMRI runs.

To provide an alternate way of testing the influence of narrative events over transitions, we correlated the number of events in each movie clip with the degree of group alignment of transitions for that clip. For each clip, we divided the number of events in that clip by its duration in minutes. For each participant, we obtained their clip conformity by taking the Fisher transformation of the correlation between the log of their step distance vector and the log of the median group signal (excluding the step distance vector of the participant in question). This was repeated for each clip within a movie run rather than the movie run in its entirety; thus, each participant had a vector describing their clip conformity, wherein each element corresponded to a conformity value for a particular clip. We calculated the Pearson correlation between the events per minute and conformity vector within each participant, resulting in a distribution of 184 correlation coefficients. We fed this distribution into a bootstrapping analysis (as described previously when we used this strategy to obtain 95% confidence intervals) with 1000 samples to test whether the correlation between events per minute and group alignment was significantly different from 0.

#### Step 3: Linking movie and rest

##### Spatial correlates of transitions versus meta-stable timepoints shared across task and rest

Using the transition and baseline onsets, we next performed a voxel-wise conjunction analysis in which we sought to identify stable spatial correlates of network meta-state transitions that could be found across rest and movie runs. Through this analysis, we wished to learn whether any set of transition predictors could link brain activity for which we have insight into psychological relevance (mv-fMRI) with brain activity for which we do not (rs-fMRI). To this end, we sampled the average fMRI image at transition and baseline onsets for each participant in the same manner as with feature vectors above, but without correcting for HRF lag (since the predictor and dependent variables were affected by the same delay). In this case, however, we created participant-wise transition and meta-stable timepoint averages not only for mv-fMRI runs, but also (separately) for rs-fMRI runs. We also spatially smoothed each image using a 6 mm FWHM gaussian kernel (at the lower bound of optimal parameters for overcoming inter-subject variability (Mikl et al., 2007).

#### Step 4: Predicting neuroticism using transition rate and conformity

The HCP administered the Neuroticism/ Extroversion/Openness Five Factory Inventory (NEO-FFI; McCrae and Costa, 2004). We used participant’s scores for each facet of human personality (neuroticism, extroversion/introversion, agreeableness, openness, and conscientiousness) as correlates to average rs-fMRI transition rate and conformity (specific to mv-fMRI). We used a bootstrap correlation approach with 1,000 samples to assess correlation between neuroticism and transition rate.

**Table S1.** Voxel clusters showing reliable activity differences at network transition vs. meta-stable timepoints in both movie-viewing and resting fMRI. Related to: Figure 4.

| Region | BA | Hemi. | Peak MNI coordinates |  |  | Peak z estimate | Spatial extent (mm³) |
| --- | --- | --- | --- | --- | --- | --- | --- |
|  |  |  | X | Y | Z |  |  |
| Transition > Meta-stable |  |  |  |  |  |  |  |
| Frontal lobe |  |  |  |  |  |  |  |
| Dorsal anterior cingulate ctx. | 32 | L/R | -3 | 13 | 37 | 6.59 | 2232 |
| Ventrolateral prefrontal ctx. | 6 | R | 56 | 0 | 11 | 5.06 | 819 |
| Insular Lobe |  |  |  |  |  |  |  |
| Anterior insula | 13 | R | 43 | 11 | -2 | 6.18 | 1282 |
| Posterior insula | 13 | R | 45 | -11 | -6 | 5.53 | 754 |
| Parietal lobe |  |  |  |  |  |  |  |
| Posterior cingulate ctx. | 23 | L/R | -5 | -38 | 24 | 7.69 | 1962 |
| Precuneus | 7 | L/R | -6 | -43 | 58 | 6.24 | 2597 |
| Visual lobe |  |  |  |  |  |  |  |
| Visual assoc. ctx. | 18 | L/R | 2 | -70 | 21 | 7.15 | 6230 |
| Meta-stable > Transition |  |  |  |  |  |  |  |
| Frontal lobe |  |  |  |  |  |  |  |
| Angular g. | 39 | R | 53 | -61 | 27 | -5.37 | 819 |
| Angular g. | 39 | L | -40 | -67 | 48 | -5.29 | 1442 |

Note: Peak coordinates are displayed in MNI space (Cocosco et al., 1997).

### Quantification and Statistical Analysis

#### Bootstrapping analyses

We used a bootstrapping approach to determine confidence intervals, descriptive statistics, and for statistical evaluations throughout the analysis.

##### Confidence intervals

We used a bootstrap approach to determine the stability of the step distance vectors across multiple t-SNE algorithm runs and to find 95% confidence intervals for eta-squared values in the psychological relevance analysis.

For the step distance vectors, we fed the step distance vectors resulting from all 100 iterations into a bootstrapping analysis. Then, for each timepoint, the 100 values from each iteration were resampled with replacement and the mean of that sample recorded. This resampling process was repeated 1,000 times to build a sampling distribution of the mean for each timepoint. Confidence intervals around each element of the participant’s mean step distance vector are derived from the 5^th^ and 95^th^ percentile values of the corresponding sampling distribution.

For the eta-squared values, we first obtained a vector for each feature describing the difference between average transition and meta-stable timepoint feature values within-participant (i.e., 184 elements representing the difference for each participant).

The transition-meta-stable difference vectors for each feature were fed into the bootstrap analysis to obtain 95% confidence intervals around the mean difference. Then, these bounds replaced the mean in the eta-squared calculations to obtain corresponding eta-squared values at the bounds of the confidence interval.

##### Descriptive statistics

We followed a similar procedure as described above to bootstrap the mean values for conformity during movie-viewing, conformity at rest, the mean correlation between events per minute and conformity (within-participant). Specifically, the input data (e.g., conformity values across participants’ movie runs) was resampled with replacement 1,000 times. The mean of each sample was recorded to build a sampling distribution of the mean, from which we could obtain the bootstrapped mean value as well as confidence intervals.

##### Correlation

We used a bootstrap approach to determine correlations between transition rate during movie-viewing and at rest, as well as between transition rate, conformity, and neuroticism. Similar to above, this entailed resampling with replacement pairs of values (e.g., movie and rest transition rates) across participants. The correlation between variables for each sample was recorded to build a sampling distribution of correlation, from which we could obtain the bootstrapped correlation value, confidence intervals, and p-values.

##### Conjunction analysis

To assess regions evoked at transition versus meta-stable timepoints in both task and rest, we first subtracted meta-stable from transition images independently for movie and rest, such that a movie difference image and rest difference image was available for each participant. We masked these images using a grey matter mask, performing comparisons only on those voxels with at least a 50% probability of being grey matter based on an MNI anatomical atlas in the same space (Mazziotta et al., 1995). Again, we used a non-parametric bootstrapping analysis, but this time obtained a bootstrap ratio *image* for each of the movie and rest difference images (see e.g., Poppenk and Norman, 2014). As the bootstrap ratio approximates a z-distribution (McIntosh and Mišić, 2013), we used a cumulative distribution function to convert it into a map of voxel-wise *p-*statistics. To perform conjunction analysis, we then thresholded both the movie and rest *p*-maps at a *P* < 0.05, setting all voxels above this value to infinity, and computed the product of the *p*-maps. The resulting conjunction *p*-map represented the probability of obtaining a supra-threshold result not only in map A, but also in map B. Because of the initial thresholding step, it had an implied *p-*value threshold of 0.0025. We then further suppressed supra-threshold voxels within the conjunction *p*-map that did not satisfy a minimum cluster extent threshold of 150 voxels (614.4 mm^3^). We selected these voxel-wise and extent thresholds to achieve a balance between Type I and Type II error rate (Lieberman and Cunningham, 2009).

### Data Availability

Data are available from the Human Connectome Project at humanconnectome.org. Movie-related variables (i.e., event boundaries that were segmented for this specific study) are available upon request.

## Supplemental Item Titles

**Movie S1. Movie-based elicitation of meta-state transitions in a single participant.**

A unique strength of the current approach is the ability to leverage sufficient signal from brain data to identify meaningful transitions in a participants’ cognitive state without any averaging across time or use of priors from another session or group. To illustrate the signal-to-noise ratio of this “stream” of cognitive state changes, we overlaid a bell sound on the movie audio of a sample participant to denote times at which meta-state transitions were elicited in that individual (with correction for the hemodynamic response function). These times were derived from peaks in the participants’ step distance vector and were derived without reference to the group (i.e., this analysis could have been performed using a single pilot participant). Meta-state transitions during the clips strongly aligned to semantic change in the film stimulus, similar to past results of brain-based movie event segmentations using averages across the entire study population (e.g., Baldassano et al., 2017). Such alignment supports the interpretation that the transitions are meaningful even at a single-subject level (see Figure 2B for systematic analysis of such variance). Related to: Figure 2B.

